# A genetic trap in yeast for inhibitors of SARS-CoV-2 main protease

**DOI:** 10.1101/2021.09.14.460411

**Authors:** Hanna Alalam, Sunniva Sigurdardóttir, Catarina Bourgard, Ievgeniia Tiukova, Ross D. King, Morten Grøtli, Per Sunnerhagen

## Abstract

The ongoing COVID-19 pandemic urges searches for antiviral agents that can block infection or ameliorate its symptoms. Using dissimilar search strategies for new antivirals will improve our overall chances of finding effective treatments. Here, we have established an experimental platform for screening of small molecule inhibitors of SARS-CoV-2 main protease in *Saccharomyces cerevisiae* cells, genetically engineered to enhance cellular uptake of small molecules in the environment. The system consists of a fusion of the *E. coli* toxin MazF and its antitoxin MazE, with insertion of a protease cleavage site in the linker peptide connecting the MazE and MazF moieties. Expression of the viral protease confers cleavage of the MazEF fusion, releasing the MazF toxin from its antitoxin, resulting in growth inhibition. In the presence of a small molecule inhibiting the protease, cleavage is blocked and the MazF toxin remains inhibited, promoting growth. The system thus allows positive selection for inhibitors. The engineered yeast strain is tagged with a fluorescent marker protein, allowing precise monitoring of its growth in the presence or absence of inhibitor. We detect an established main protease inhibitor down to 10 μM by a robust growth increase. The system is suitable for robotized large-scale screens. It allows *in vivo* evaluation of drug candidates, and is rapidly adaptable for new variants of the protease with deviant site specificities.

**IMPORTANCE:** The COVID-19 pandemic may continue several years before vaccination campaigns can put an end it globally. Thus, the need for discovery of new antiviral drug candidates will remain. We have engineered a system in yeast cells for detection of small molecule inhibitors of one attractive drug target of SARS-CoV-2, its main protease which is required for viral replication. To detect inhibitors in live cells brings the advantage that only compounds capable of entering the cell and remain stable there, will score in the system. Moreover, by its design in yeast, the system is rapidly adaptable for tuning of detection level, eventual modification of protease cleavage site in case of future mutant variants of the SARS-CoV-2 main protease, or even for other proteases.

## INTRODUCTION

The COVID-19 pandemic caused by SARS-CoV-2 is currently ongoing since a year and a half. Despite unparalleled successes in vaccine development and roll-out of vaccination programs, it is uncertain when this will be sufficient to quell the outbreak, in the face of arising mutant virus strains. For the foreseeable future, the need for antiviral drugs and treatment for COVID-19 remains. The initial attempts at drug discovery have been dominated by repurposing of antiviral and other drugs. Previous experience of discovery and development of antiviral drugs against coronaviruses is limited. The SARS epidemic in 2003, and MERS in 2012, did spur some early drug discovery efforts, *e*.*g*. (1-3). However, none of those have been further developed into clinically approved antiviral drugs. Given the similarity between the coronaviruses MERS-CoV, SARS-CoV-1, and SARS-CoV-2, it is likely that an antiviral drug effective against any of the first two could also be effective against SARS-CoV-2. This argues that continued research through multiple avenues on new antiviral drug candidates targeting SARS-CoV-2 will be warranted for the next years.

The coronavirus main protease (Mpro; also known as poliovirus 3C-like protease, 3CL pro) cleaves the viral polyprotein at 11 sites into its individual functional protein products, and is essential for the virus to replicate. It is recognized as a suitable target by virtue of its critical role in viral propagation, and by the well explored druggability of proteases. There are no human host-cell proteases structurally related to SARS-CoV-2 Mpro and no human protease shares its strong preference for cleaving C-terminally of a glutamine. By contrast, the SARS-CoV-2 papain-like protease – also an antiviral target candidate – is expected to have overlapping substrate and inhibitor sensitivity profile with host cell deubiquitinases (4).

The crystal structure of Mpro has been determined (5), opening up for structure-based drug discovery. However, it has been argued that the dynamic properties of SARS-CoV-2 Mpro, in particular through flexible regions near the catalytic active site, make it a more difficult target for rational drug discovery than the SARS-CoV-1 Mpro (6).

An *in vivo* assay of candidate inhibitors designed to act intracellularly confers the advantage that only molecules able to be efficiently taken up by cells and remain stable in the intracellular environment will score in the assay. Testing of candidate molecules in assays with purified enzyme *in vitro* is sensitive to buffer conditions, such as recently demonstrated for putative SARS-CoV-2 Mpro inhibitors tested in a variety of *in vitro* assays, where the apparent inhibition was obliterated by restoration of a reducing environment (7). Using yeast (*S. cerevisiae*) gives access to a versatile platform for precise and rapid genetic engineering of intracellular testing systems. Genetic screens in yeast have previously identified small molecule inhibitors to *e*.*g*. HIV-1 protease (8), SARS-CoV-1 papain-like protease (9), and to SARS-CoV-1 mRNA cap-methyltransferase (10).

Here, we have constructed a yeast strain expressing a synthetic fusion protein, which upon cleavage by a protease releases a bacterial toxin moiety capable of inhibiting growth of the yeast strain. In the same strain, SARS-CoV-2 Mpro is expressed, and its recognition site is inserted in the fusion protein, making cell growth negatively regulated by the Mpro activity level, but stimulated by Mpro inhibitors (Fig. 1). The yeast strain genetic background is sensitized to external molecules by inactivation of membrane-bound transporters. We demonstrate detection of an established Mpro inhibitor by growth promotion of engineered yeast cells, in concentrations down to 10 μM. The system is suited for physical screening in robotized setups that allow controlled growth of yeast cells in microtiter format and measurement of fluorescence intensity. The design enables rapid upgrades of the protease recognition site according to eventual future mutant variants of the virus.

**Figure 1.**
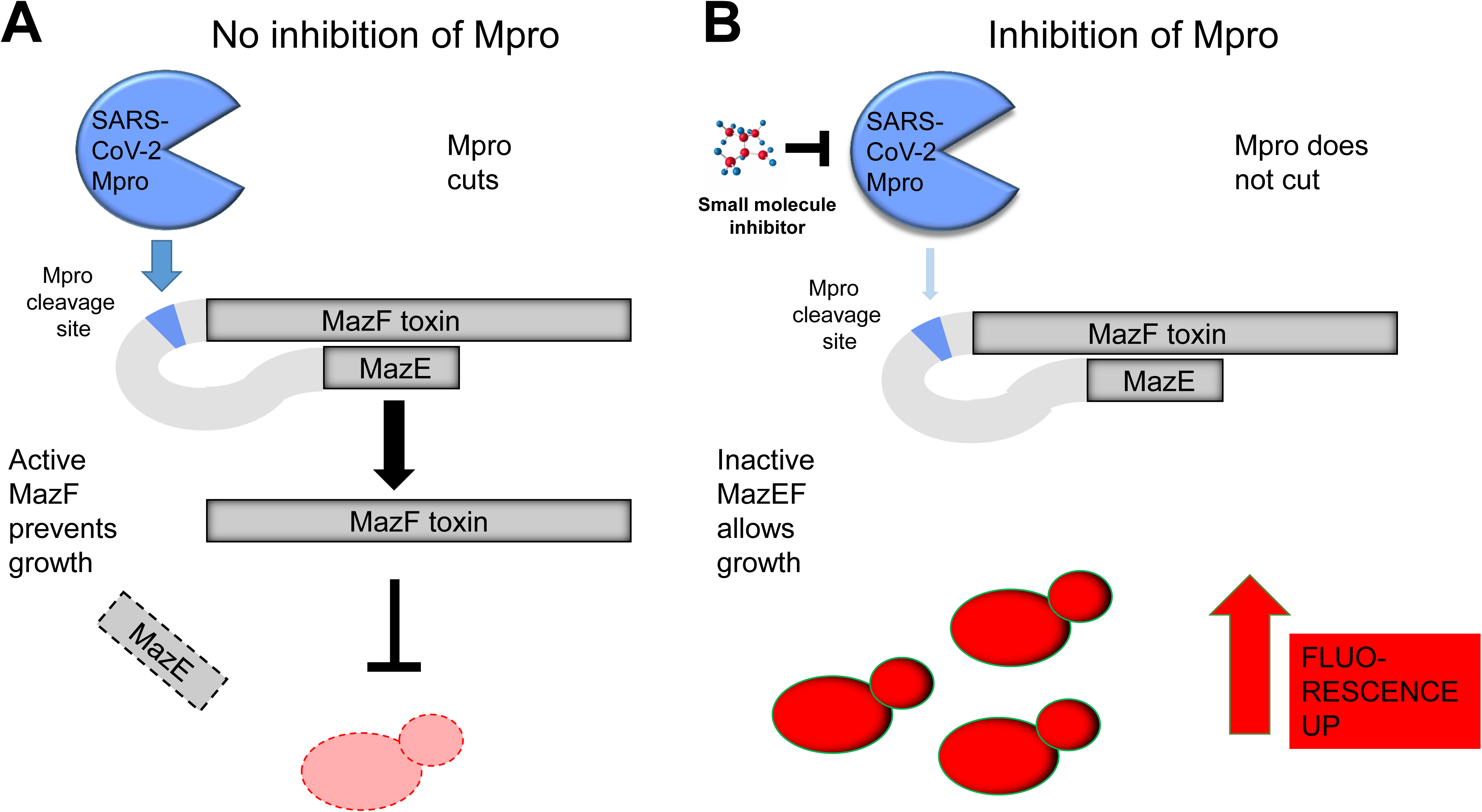
Principle of the genetic selection system **A)** Situation without Mpro inhibitor. SARS-CoV-2 Mpro is active, and cuts the inactive MazEF fusion protein at the synthetic Mpro cleavage site in the linker region connecting the MazF (toxin) and MazE (toxin inhibitor) moieties, releasing active MazF. Then, the RNase activity of MazF can exert its toxic effect by degrading cellular RNAs, thus preventing growth of the cells carrying the construct. The output signal from the red fluorescent protein tag in these cells will be weak. **B)** Situation with Mpro inhibitor. The activity of SARS-CoV-Mpro is reduced, and the inactive MazEF fusion stays mostly intact. The toxic RNase activity will be reduced, and these cells can grow faster, resulting in a stronger fluorescent signal output.

## RESULTS

### Construction of a genetic trap in yeast

The *E. coli* toxin MazF is a single-strand endoribonuclease which cleaves mRNA at ACA sequences (11). When expressed in either a prokaryotic or eukaryotic cell, it arrests growth or kills that cell, depending on the expression level. MazF is a component of a chromosomally encoded toxin/antitoxin system in *E. coli*, the other component being MazE, a protein binding to and inhibiting MazF. MazE has a much shorter turnover time than MazF, requiring the cell to continuously synthesize MazE to maintain viability (12).

It has previously been demonstrated that a synthetic protein fusion of a C-terminal fragment of MazE to MazF resulted in a chimeric protein with inhibition of MazF activity, permitting cell survival and growth also at high expression levels (13). Insertion of a protease site in the added linker sequence connecting the two protein moieties permi release of fully active MazF upon expression of the protease and cleavage of the linker peptide (13). To exploit these findings for creation of a genetic selection system in *S. cerevisiae*, we generated a chimeric construct (MazEF) with the Mpro cleavage site in the linker. Specifically, the construct contained the 41 C-terminal amino acid residues (aa) Leu42 to Trp81 of MazE with an added ATG codon preceding the Leu42 codon, followed by a GGVKLQSGS aa linker sequence containing the Mpro cleavage site VKLQS, joint N-terminally to the full aa sequence of MazF. The entire coding region was modified using silent mutations making it devoid of ACA sequences to reduce cleavage of *MazF* mRNA. The coding sequence of this fusion was put under transcriptional control of the *MET3* promoter in a pCM188 (14) backbone or under the control of the *GAL1* promoter in the same backbone in order to achieve two ranges of expression; weak and strong, respectively (Table 1).

**Table 1.**
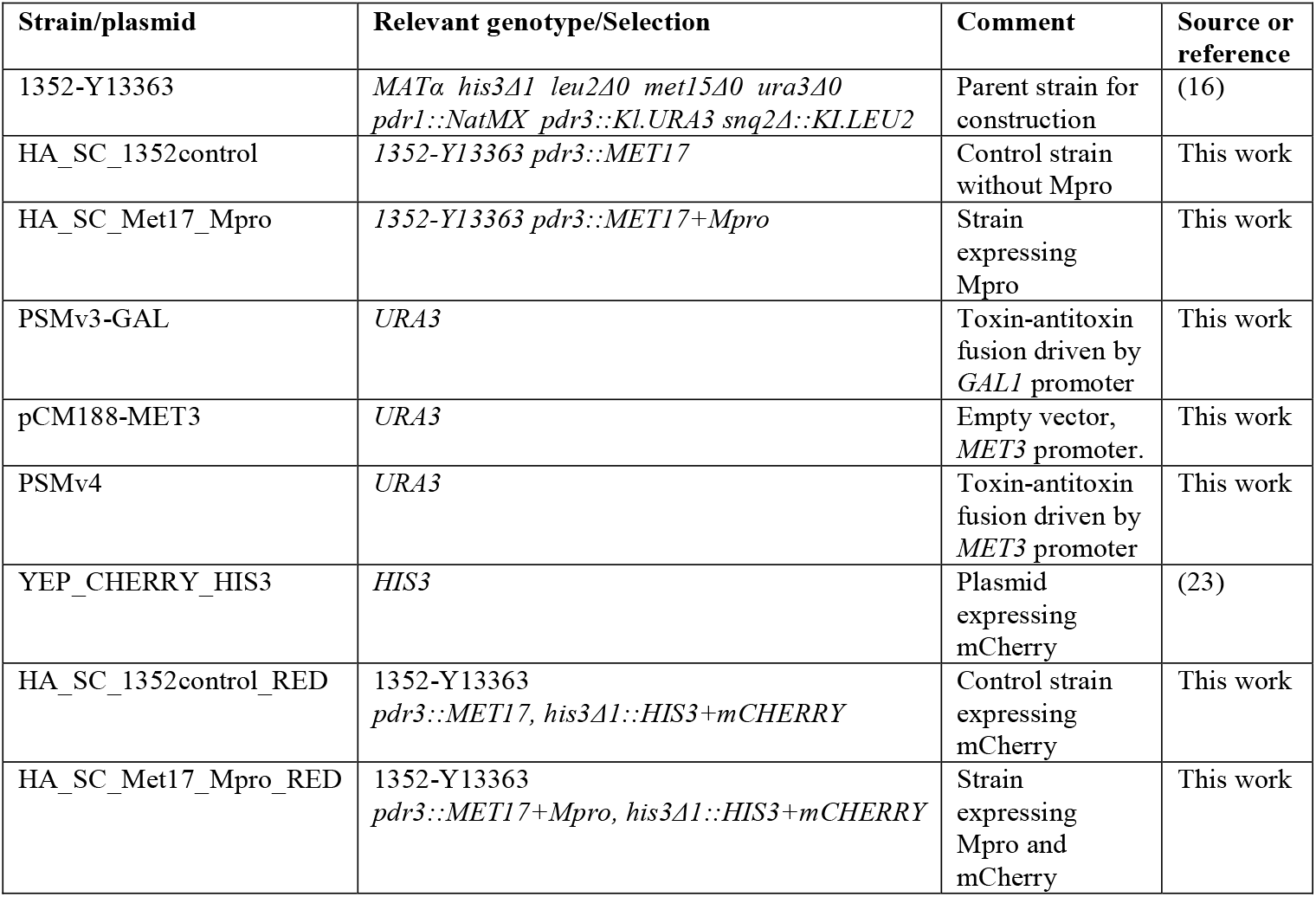
Strains and plasmids used in this study.

The genetic background for strain constructions was *S. cerevisiae* strain 1352-Y13363 (Table 1). It carries null alleles of *PDR1* and *PDR3*, encoding transcriptional regulators of a large number of ABC transporters exporting small molecules from the cell, and of *SNQ2*, which encodes an ABC exporter protein important for detoxification (15). This triple mutation greatly increases the number of compounds to which yeast cells are sensitive, without compromising vigor (16). In this strain, the *Kluyveromyces lactis URA3* marker disrupting the *PDR3* locus was first replaced by *S. cerevisiae MET17*, to create the control strain and allow methionine repressible expression from the *MET3* promoter (17) by restoring methionine prototrophy. The Mpro expressing strain was constructed by ligating a synthetic fragment carrying Mpro under the control of the *Pichia GAP* promoter with *MET17* and integrating in the *PDR3* locus (Supplementary File S1). The strains were tagged with the red-fluorescent mCherry marker under control of the constitutive *TDH3* promoter, which was integrated into the *HIS3* locus. Full details of strain construction are given in Supplementary File S1.

### Responsiveness of engineered strains to the GC376 protease inhibitor

We wanted to evaluate the engineered yeast strains for their usefulness as a testing platform for potential Mpro inhibitors. To do this, we tested the impact of expressing MazEF at different levels in the presence of Mpro, and of externally added small molecules.

GC376 is a broad-spectrum antiviral molecule active against 3CL-like proteases from coronaviruses (18) which has been used to treat feline infectious peritonitis (19). GC376 is a prodrug, and was recently shown to be effective against SARS-CoV-2 Mpro *in vitro* (20, 21) and against SARS-CoV-2 in human cell culture (21, 22). We chose this compound as a positive control for an Mpro inhibitor to titrate the growth response.

We first investigated a strain expressing the MazEF chimera from the *GAL1* promoter and Mpro from the *Pichia GAP* promoter (Supplementary File S1). However, the addition of CG376 even at 100 μM only marginally improved growth in inducing galactose medium (Supplementary Fig. S1). We concluded that expression of the MazEF toxin fusion protein from the strong *GAL1* promoter in the presence of Mpro was too detrimental to be overcome by protease inhibitors. Thus, this expression system is impractical for screening purposes.

Reasoning that we needed weaker and regulatable expression of the toxin chimera, we next examined the behavior of strains expressing Mpro from the *Pichia GAP* promoter and the toxin chimera from the *MET3* promoter, in varying methionine concentrations. As seen in Fig. 2 A, growth of the strain expressing both Mpro and MazEF is increasingly depressed by lower methionine concentrations in the range below 350 μM. This is expected since *MET3* is negatively regulated by methionine (17). By contrast, growth of strains expressing only Mpro, only MazEF, or neither, is not significantly affected by the methionine concentration (Supplementary Fig. S2, left column). This indicates that the negative impact of expression of Mpro alone in yeast cells is negligible under these conditions, and that MazEF is inactive and not cleaved to release active MazF in the absence of Mpro, in line with the results with MazEF expression from the *GAL1* promoter (Supplementary Fig. S1).

**Figure 2.**
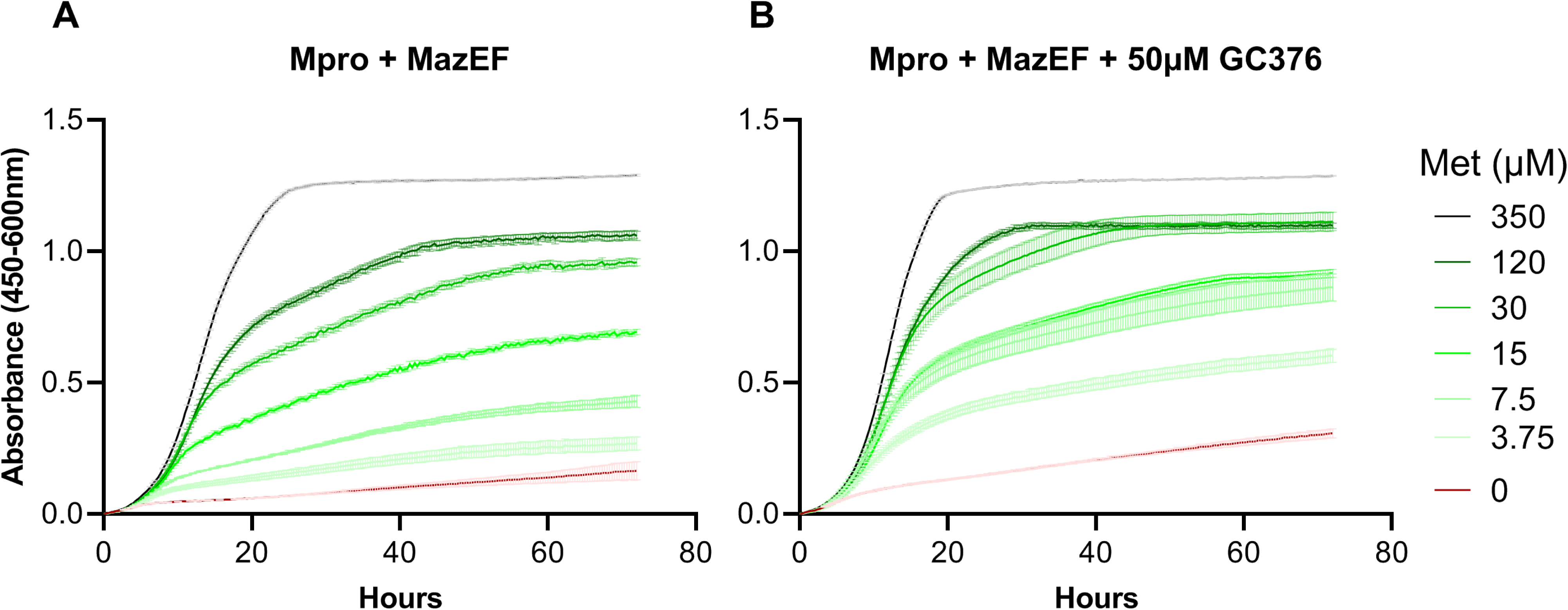
Titration of methionine concentration with yeast strain expressing Mpro and MazEF chimera (strain HA_SC_Met17_Mpro carrying PSMv4) in presence or absence of GC376 Growth measurements were obtained using the Bioscreen C reader, starting absorbance adjusted to 0 (mean ± SE, n = 3). **A)** Cell cultures were grown in SD medium without uracil (SD-ura) and with varying concentrations of methionine from 350 µM to 0 µM (see legend) **B)** The same conditions but treated with the protease inhibitor GC376 (50 µM).

To achieve maximal separation between cells exposed to an Mpro inhibitor, we then grew cells expressing MazEF and Mpro in increasing methionine concentrations up to 350 μM and compared growth with or without GC376 at 50 μM (Fig. 2 B). As expected if GC376 inhibits Mpro and therefore prevents cleavage of MazEF, releasing toxic MazF, growth was improved by the presence of GC376. The best absolute separation was seen at 7.5 - 15 μM methionine. To verify that the improved growth was indeed due to decreased Mpro activity, we then exposed cells to a range of GC376 concentrations up to 100 μM, maintaining a fixed methionine concentration of 7.5 μM. GC376 at 50 μM did enhance growth of the strain expressing both Mpro and MazEF (Figs. 2 B and 3 A), but as expected had no effect on growth of any of the strains lacking Mpro, MazEF, or both (Supplementary Figs. S2, S3). Finally, we wanted to see if lower inhibitor concentrations than 50 μM could also be detected with strains carrying the mCherry fluorescent marker. With methionine at 10 μM or lower, we could discriminate the effect of GC376 down to 10 μM (yield ratio = 1.57, SE ± 0.02, n = 16, p-value < 2.2e-16; see Supplementary Fig. S4). This also demonstrates that tagging cells with mCherry did not cause a noticeable change in their response to methionine or GC376 concentration (Fig. 3, Supplementary Fig. S3).

**Figure 3.**
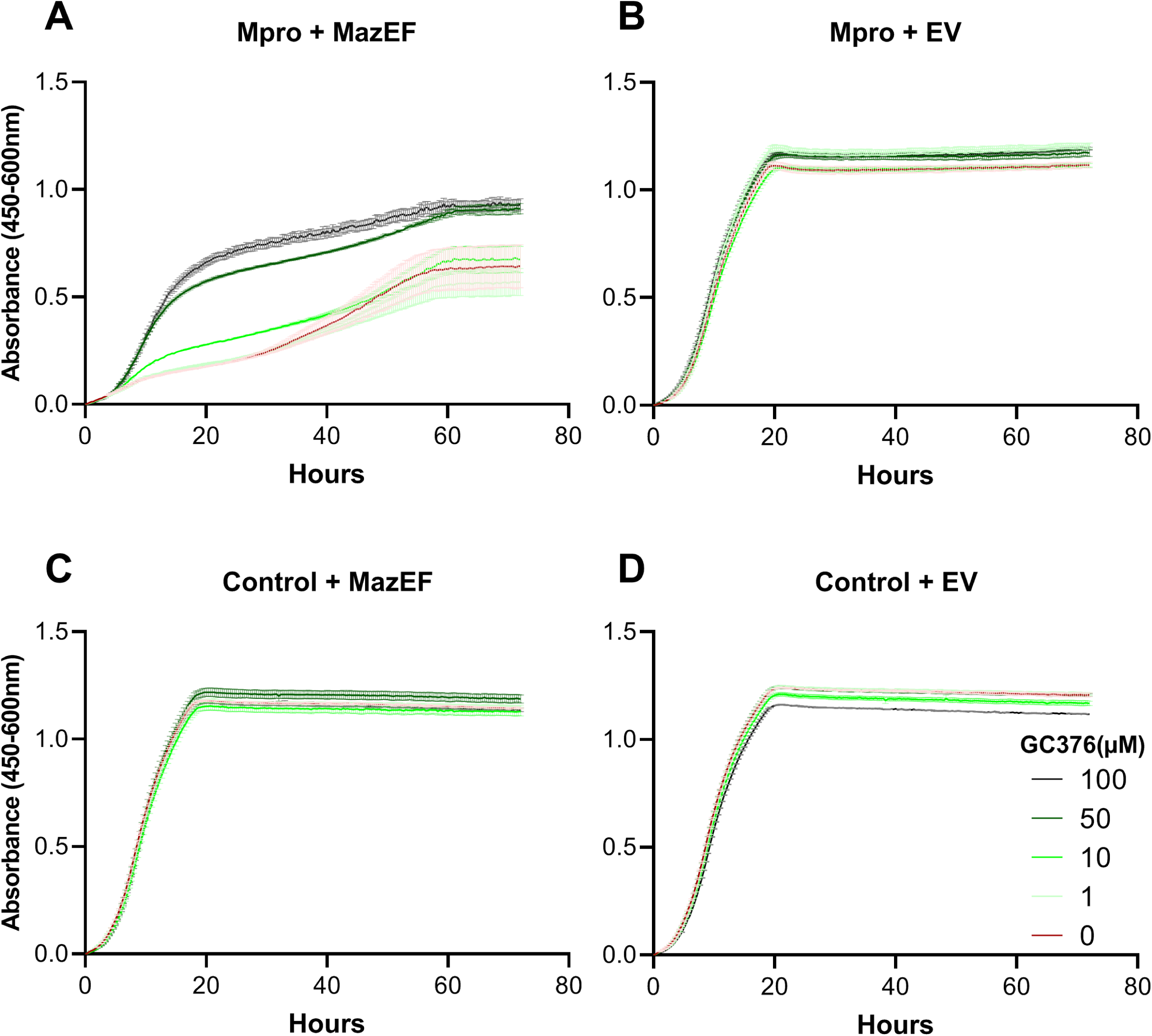
Titration of inhibitor GC376 Growth measurements were obtained as in Figure 2 (mean ± SE, n = 3). Cell cultures were grown in SD -ura and 7.5 µM methionine and varying concentrations of GC376 (see legend). **A)** Strain HA_SC_Met17_Mpro carrying PSMv4 expressing Mpro and MazEF chimera **B)** Strain HA_SC_Met17_Mpro expressing Mpro (EV; empty vector) **C)** Strain HA_SC_1352control carrying PSMv4 expressing the MazEF chimera only **D)** Control strain HA_SC_1352control with empty vector

### Measuring impact of candidate protease inhibitors on strains using a fluorescence readout

Having tuned the expression levels of the MazEF chimera from the *MET3* promoter to achieve a robust impact on growth by the inhibitor GC376, we wanted to test the screening system in a robotized platform suitable for screening of large compound libraries. Quantifying growth as fluorescence of a genetically tagged marker protein potentially offers better signal to noise ratios than optical density. Thus, we monitored growth of a strain expressing MazEF and Mpro, and in addition tagged with mCherry, in a robotic platform allowing automated growth monitoring of thousands of samples simultaneously and regular fluorescence readings (23).

Besides GC376, we exposed the strain to other compounds implicated by different criteria as potential Mpro inhibitors. Boceprevir is a clinically approved human hepatitis C virus N53 protease inhibitor, showing activity against SARS-CoV-2 Mpro in cultured human cells (22). TDZD-8, originally characterized as an inhibitor of glycogen synthase kinase β (GSK3-β), scored as a SARS-CoV-2 Mpro inhibitor in an *in vitro* assay, but was subsequently dismissed as a false positive because of its aggregation properties (24). Efonidipine and Lercanidipine are Ca^2+^ channel blockers found as hits in an *in silico* screen for SARS-CoV-2 Mpro inhibitors and also active in an *in vitro* enzymatic assay (25). Glycyrrhizic acid at high concentrations blocks SARS-CoV-2 replication in cell culture and inhibits Mpro in an *in vitro* assay (26). Compounds were applied at 10 and 30 μM. We adjusted the methionine concentration to 15 μM to avoid a gradual increase in toxicity caused by methionine consumption towards the end of the run, as can be seen in Fig. 3 A at 10 μM concentration.

In Fig. 4 A we see that based on fluorescence readings in this setting, the growth-promoting effect of CG376 on the tester strain is dose-dependent and obvious also at 10 μM. This demonstrates that the system works well also in a setting with limited aeration and fluorescence-based monitoring of growth. For the other compounds, no positive effect is seen at any concentration. TDZD-8 instead displayed a negative impact on growth (Fig. 4 H). The rest of the compounds tested; Boceprevir, Efonidipine, glycyrrhic acid, and Lercanidipine, did not impact growth (Fig. 4). Readings of optical density, which were obtained simultaneously, closely paralleled the fluorescence values. As seen in Table 2, the discrimination for the tester strain between treatment with GC376 and no treatment was best for fluorescence readings at 20 h both for 10 and 30 μM of inhibitor (ratio 1.59 or 2.28, respectively). This is better than what was achieved with reading optical density (ratio 1.22 or 1.61).

**Figure 4.**
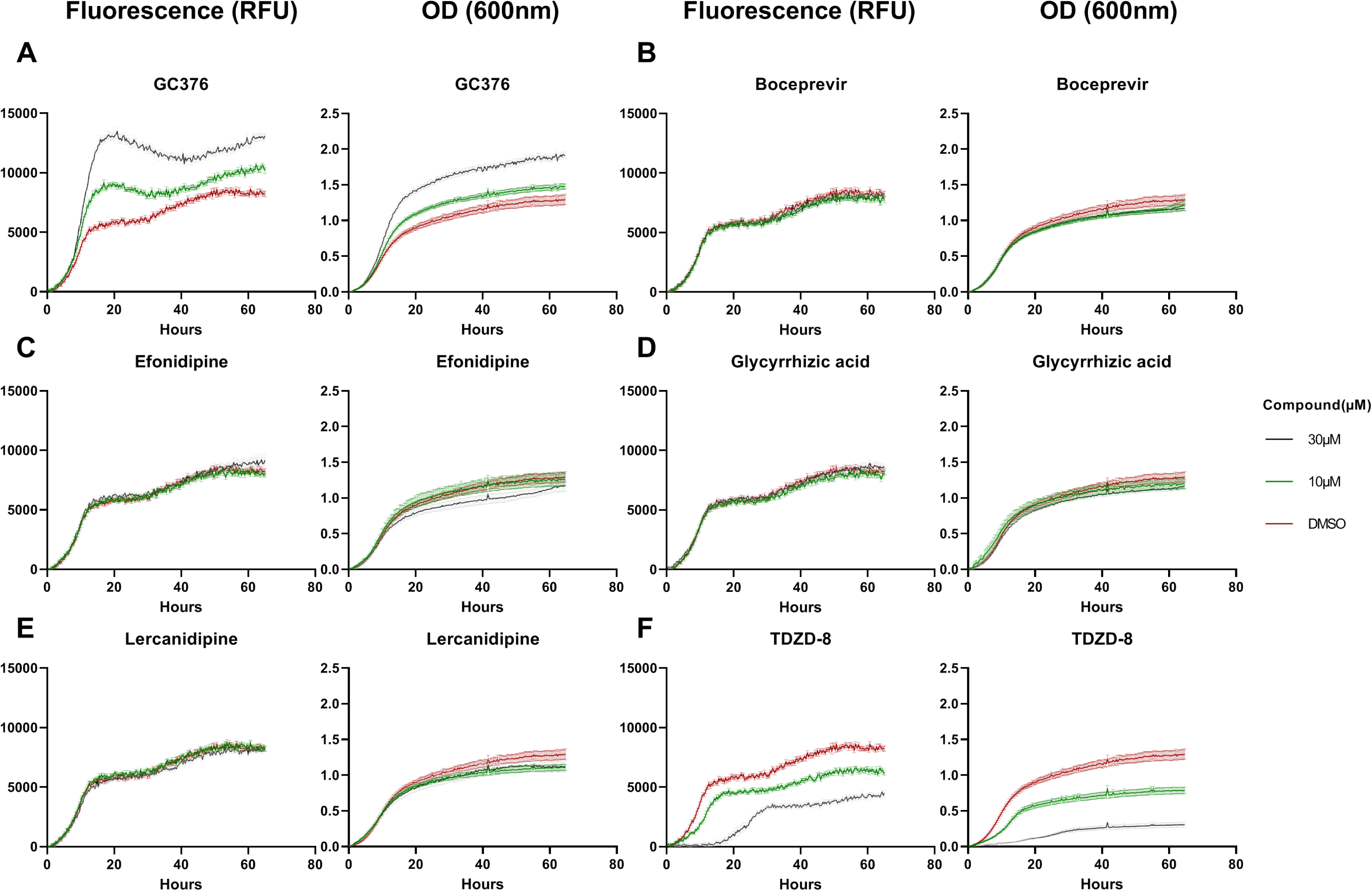
Growth of mCherry-tagged strain expressing Mpro and MazEF chimera in presence of different candidate protease inhibitors Strain HA_SC_Met17_Mpro_RED carrying PSMv4 was used. Growth measurements (fluorescence and absorbance) were obtained using the Eve robot (see Methods), starting measurement adjusted to 0 (mean ± SE, n = 16). RFU, relative fluorescence units. Cell cultures were grown in SD -uracil and 15 µM methionine; 1.25 % DMSO; and 0, 10 and 30 µM of the respective compound to be tested (see legend).

**Table 2.**
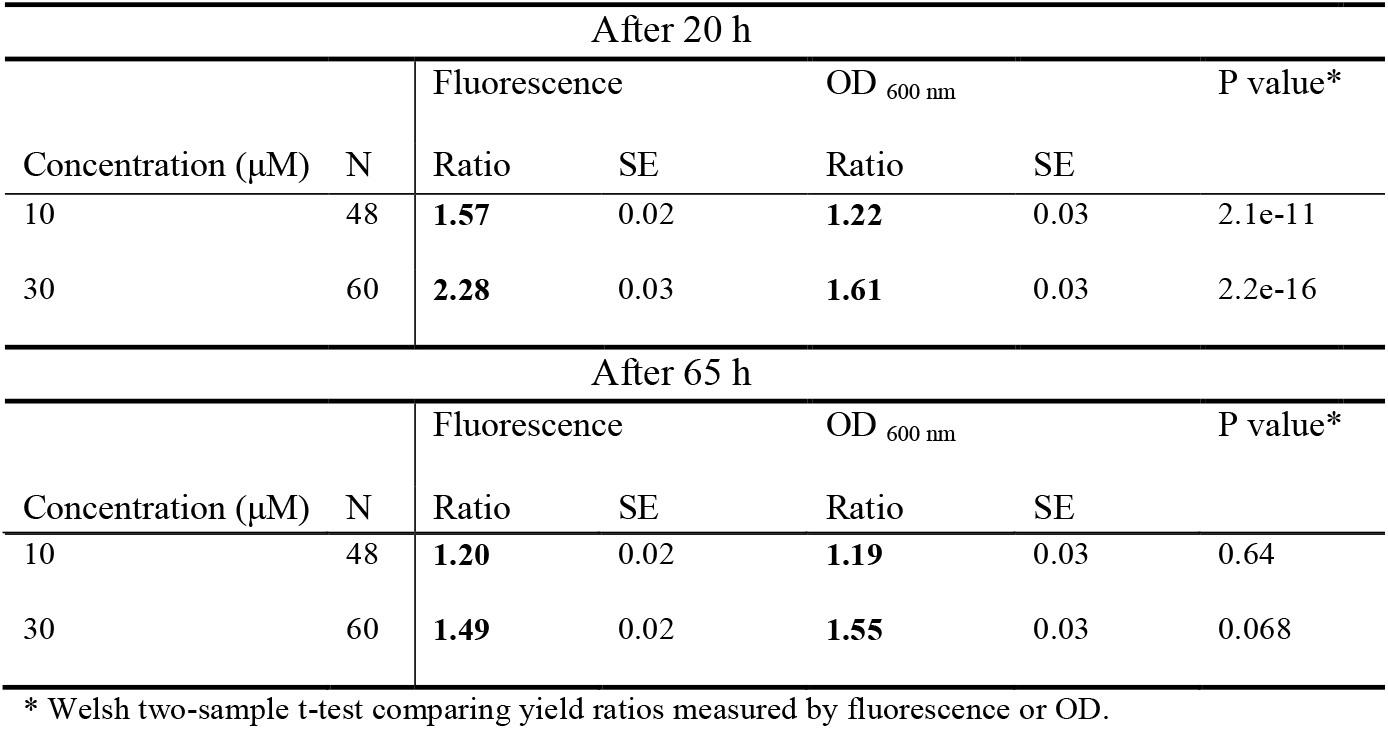
Signal discrimination for detection of Mpro inhibitor (GC376) Ratio of yield compared to controls without inhibitor

## DISCUSSION

We have created an experimental system for physical screening of candidate inhibitors of SARS-CoV-2 Mpro in yeast cells. We demonstrate that it can be used to detect with good confidence a known Mpro inhibitor at 10 μM. The growth of control strains lacking MazEF, Mpro, or both, is unaffected by the inhibitor, showing the specificity of the assay. The system is suitable for screening of chemical libraries in a robot platform capable of cultivating large numbers of yeast cells in controlled conditions with regular monitoring of fluorescence. Using a fluorescent readout provides increased sensitivity over optical density reading, as the background signal is decreased, and issues such as cell clumping or inhomogeneities in the culture medium do not affect the output signal.

The dynamics of the effects on growth in this system are determined by the two players, MazEF and Mpro. At the onset of an experimental cycle, *MazEF* expression is kept low by the high methionine concentration in the preculture medium, while both Mpro and mCherry are expressed at high constitutive levels. On transfer to the culture medium, the external methionine concentration drops, and transcription of *MazEF* from the *MET3* promoter starts to rise. The toxic effect of active MazF is to degrade cellular RNAs. To minimize any complications from MazF cleaving its own mRNA, we engineered the coding region of *MazF* to lack ACA sequences. In the mRNA encoding Mpro, ACA sequences were left intact, however. By expressing Mpro from the strong constitutive *Pichia GAP* promoter, we ensure that enough protease is present at the time MazEF production starts to increase through the gradual reduction of methionine in the medium from the experiment onset. Beside its intended target in the MazEF synthetic linker, Mpro can cleave endogenous yeast proteins. The consensus protease cleavage site profile of SARS-CoV-2 Mpro comprises the sequence [A/S/T]-LQ[A/S/G] (4, 27), and there are 136 potential Mpro cleavage sites in the yeast proteome (Supplementary Table S3). There is a slight negative impact of expressing Mpro alone in yeast cells (Supplementary Fig. S5), as expected from cleavage of cellular proteins. However, expression of MazEF had no effect on growth in the absence of Mpro (Supplementary Figs. S2, S3). This indicates that there is no background cleavage of MazEF by yeast proteases.

We also observed that starting the experiment at low culture density (OD_600nm_ < 0.1), such as presented here, works well for detecting Mpro inhibitors. A higher density of the preculture results in reduced sensitivity to the inhibitor, presumably because not enough MazF accumulates before the culture approaches stationary phase for it to significantly affect growth.

In the present system, GC376 gave a strong and specific signal as an Mpro inhibitor. Other molecules previously reported by various criteria to be Mpro inhibitors did not score. There are several possible explanations for this. First, the previous evidence from *e*.*g*. an *in vitro* assay could be misleading, as exemplified by aggregation-prone TDZD-8 (20). Another reason why a small molecule inhibitor may fail to show as a hit in this assay is insufficient cellular uptake. Yeast cells have multiple membrane-bound proteins, notably ABC transporters, which extrude exogenous molecules from the cell. Several attempts have been made to diminish this effect of transporters in order to sensitize yeast to small molecules, a key issue for chemical genomics and drug screening in yeast (28). A strain lacking all 16 ABC transporters has been generated (29), and a nine-tuple deletion strain lacking genes encoding transporters and transcription factors controlling similar genes displayed increased sensitivity to various compounds in the range 2 – 200 fold (15). However, this strain has poor vigor, and it was later found that most of its sensitivity can be recovered by retaining only the *pdr1Δ pdr3Δ snq2Δ* gene deletions, which leaves the strain with robust growth properties (16). Here, we use this triple deletion strain background, which should provide a higher sensitivity to small molecules and a higher scoring rate than in previous screens using the same physical cultivation and reading system and the same fluorescent marker, where only the *pdr5Δ* deletion was used to sensitize the yeast tester strain (23, 30, 31). Nevertheless, many compounds will not penetrate into the yeast cells and fail to score for that reason. Finally, inhibitors with general toxicity will not score in this assay system as the depression of growth will cancel out the positive signal from decreased protease activity. On the other hand, general toxicity is obviously to be avoided in this context. The negative impact by TDZD-8 (Fig. 4 F) could be due to general toxicity, or to the compound inhibiting the yeast homologs of GSK3-β; Rim11, Mck1, Mrk1, and Ygk3 (32).

The setup is suitable for settings with robotic handling of samples at microtiter plate scale. The ability to continuously read fluorescence and optical density of cultures during an entire growth cycle, with the ability to oxygenate the culture wells by shaking, is important to obtain reliable results. In a longer perspective, this setup may be modified to target other proteases, by exchanging the protease cleavage site in the linker of the fusion protein. The concept is general but has some constraints. First, the consensus cleavage site of the protease has to been known with sufficient accuracy. Second, the protease has to be capable of acting in an environment where the fusion protein is accessible to attack, *e*.*g*. not in the interior channel of a proteasome.

In summary, we describe a functional *in vivo* screening system capable of identifying candidate inhibitors of SARS-CoV-2 Mpro. Being a cellular assay, it selects for molecules with high bioavailability while avoiding some pitfalls of *in vitro* environments. At the same time, it is performed in a safe laboratory setting, without virus particles. It is versatile in that it can be adapted to target other proteases, benefitting from the facile genetic engineering of *S. cerevisiae*. The system has been tuned to detect an inhibitor down to 10 μM using a positive selection mode. Importantly, this avoids the false positives in a negative selection mode, where compounds with general toxicity would score as hits.

## MATERIALS AND METHODS

### Construction of plasmids and yeast strains

Details of the construction of each strain and plasmid are in Supplementary File S1. Maps of constructs are in Supplementary Files S2 – S4.

### Candidate protease inhibitor compounds

Boceprevir, GC376, TDZD-8, Lercanidipine hydrochloride, Efonidipine hydrochloride monoethanolate, and glycyrrhizic acid ammonium salt from *Glycyrrhiza* root were purchased from Sigma. Stocks of all the compounds were made in DMSO at a concentration of 20 mM and stored frozen at −80°C.

### Culture conditions

*S. cerevisiae* strains and plasmids used are listed in Table 1. YPD (2 % glucose, 2 % peptone and 1 % yeast extract) was used for routine culturing of strains not carrying plasmids. Synthetic Defined (SD) medium (0.19 % yeast nitrogen base, 0.5 % ammonium sulfate, 2 % glucose and 0.077 % Complete Supplement Mixture [CSM, ForMedium]) with appropriate drop out used for routine culturing of strains carrying plasmids and for Bioscreen C (Labsystems Oy) phenotyping (uracil dropout with varying concentrations of methionine). Liquid cultures were grown in a rotary shaker at 30°C at 200 rpm.

### Phenotypic analysis in Bioscreen

Unless stated otherwise, overnight cultures were made in SD –ura containing 350 µM methionine and maintained at exponential growth phase. Before the start of experiment, the media was removed and the pellet resuspended in SD –ura –met to a final OD of 1.0. Subsequently the cultures were diluted to an OD_600 nm_ of 0.1 into 100-well honeycomb plates (Labsystems Oy) pre-aliquoted with the appropriate media using an Opentrons OT2 robot to carry out both the dilution and media dispensing. Strains were cultivated for three days with the low shaking setting at 30°C with 20 min measurement intervals using a wide band filter (450 – 600 nm).

### Phenotypic analysis in robot Eve

Overnight cultures were grown and maintained as described above. Before the start of experiment, media was removed and the pellet resuspended in SD –ura containing 15 µM methionine to an OD_600 nm_ of 0.1. Within the automated workstation, the culture was aliquoted into a Greiner 384-well black plate with clear bottom using the Thermo Combi multidrop, and chemical compounds were transferred to the assay plate using the Bravo Liquid Handling platform to a final concentration of 10 or 30 μM of each compound (final DMSO concentration 1.25 %) and a final volume of 80 μl. Cells were incubated and their growth monitored essentially as described (23). Growth was stationary at 30°C, except at each 20-min interval, samples were circularly agitated at 1000 rpm for 10 s, followed by 10 s in the opposite direction. A read-and-incubate cycle was performed using the Overlord automation system to determine growth at 20 min intervals for 64 h. Fluorescence (580 nm excitation / 612 nm emission) and optical density (OD_600 nm_) measurements were obtained with a BMG Polarstar plate reader.

## ACKNOWLEDGEMENTS

This work was supported by a grant from the Swedish Research Council (2020-05738).

## SUPPLEMENTARY MATERIAL

**Supplementary Figure S1** Growth properties of the strain carrying Mpro with *GAL1-*driven expression of MazEF. Growth measurements were obtained using the Bioscreen C reader, starting absorbance adjusted to 0 (mean ± SE, n = 3). Cells cultures were grown in different carbohydrate sources to allow (raffinose + galactose) or neither repress nor induce (raffinose) expression of the MazEF toxin, and 100 µM GC376 or 0.2 % DMSO (see legend).

**Supplementary Figure S2** Growth of control strains expressing Mpro only (A), the MazEF chimera only (B) or neither (C) in varying concentrations of methionine.

Left panels: Cell cultures were grown in SD medium without uracil (SD-ura) and with concentrations of methionine varying from 350 µM to 0 µM (see legend). Right panels: The same conditions but treated with the protease inhibitor GC376 (50 µM).

**Supplementary Figure S3** Growth of control strains carrying mCherry tag; not expressing MazEF, not expressing Mpro, or neither MazEF nor Mpro

Growth of mCherry tagged control strains expressing Mpro only (**A**) the MazEF chimera only (**B**) or neither (**C**) in varying concentrations of methionine and GC376.

Left panels: Cell cultures grown in SD medium without uracil (SD-ura) and with varying concentrations of methionine from 350 µM to 0 µM (see legend). Middle panels: The same conditions treated with 10 µM GC376. Right panels: The same conditions but treated with 30 µM GC376.

**Supplementary Figure S4** Strain tagged with mCherry and expressing Mpro and the MazEF chimera was exposed to different GC376 and methionine (Met) concentrations.

Strain HA_SC_Met17_Mpro_RED carrying PSMv4 was used. Growth measurements were obtained as in Fig. 2 (mean ± SE, n = 3). Cell cultures were grown in SD –uracil with varying concentrations of GC376 (see legend).

**A**) 0 µM Met; **B**) 10 µM Met; **C**) 120 µM Met

**Supplementary Figure S5** Effect on yeast by expressing only MPro Cultures were grown in SD-ura (mean ± SE; n = 3).

**Supplementary File S1** Strain and plasmid construction

**Supplementary File S2** Complete sequence PSMv3-Gal

**Supplementary File S3** Complete sequence pCM188-MET3

**Supplementary File S4** Complete sequence PSMv4

**Supplementary Table S1.**
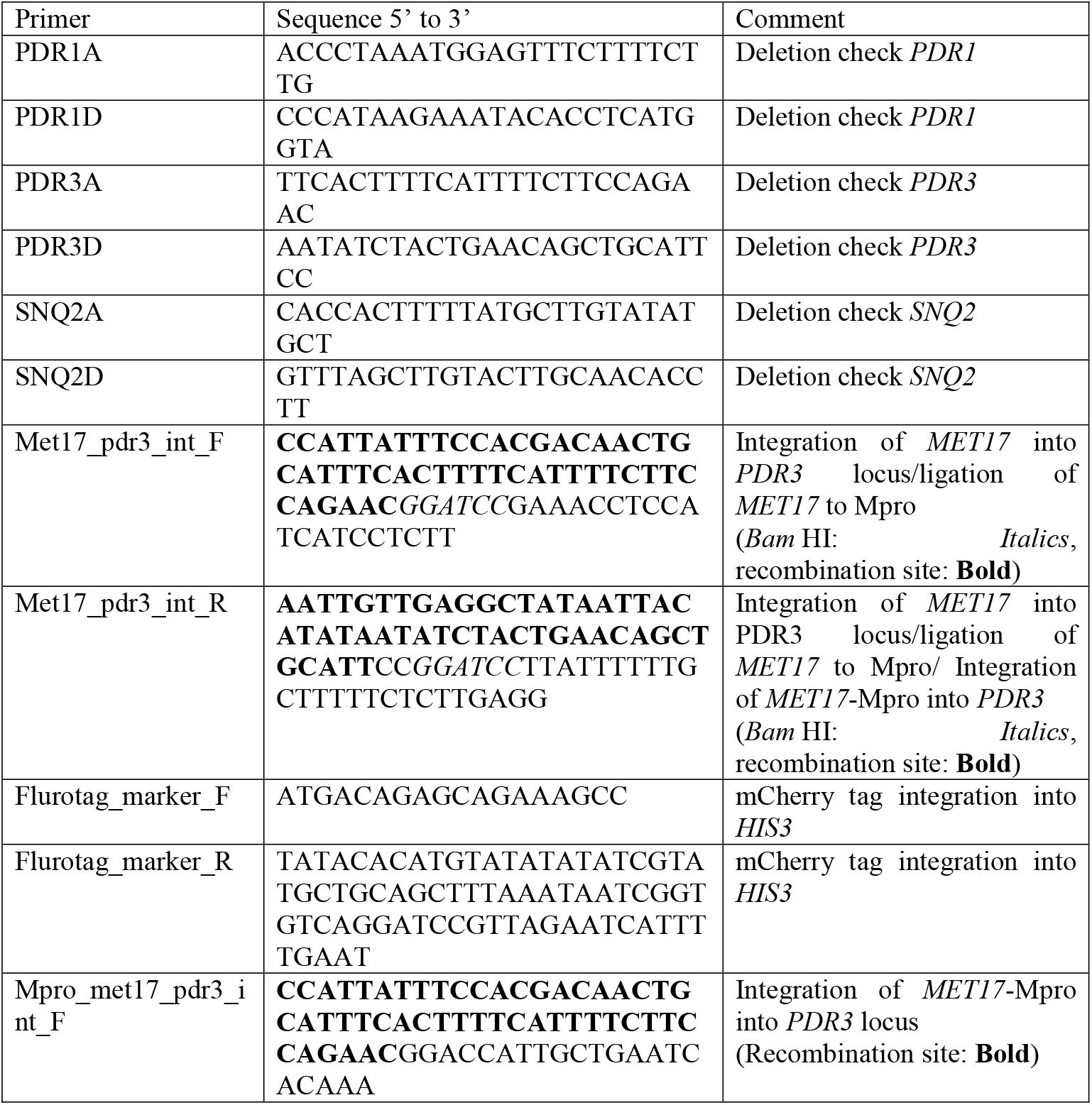
Primers used for strain construction

**Supplementary Table S2.**
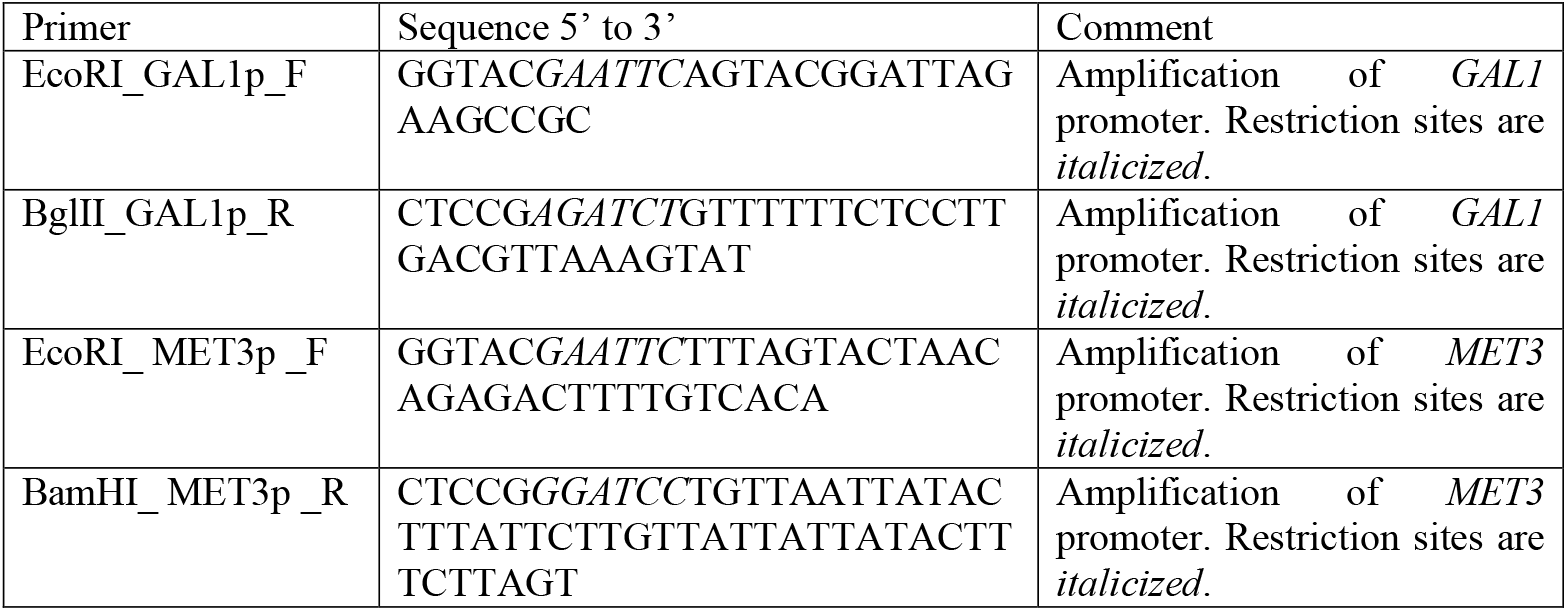
Primers used for plasmid constructions

**Supplementary Table S3** Predicted Mpro cleavage sites in the *S. cerevisiae* proteome

## Supplementary File S1

### Strain construction

#### HA_SC_1352control

Deletions of *PDR1, PDR3* and *SNQ2* in the parental strain 1352-Y13363 were confirmed by PCR. *MET17* was amplified from the CENPK2 background using primers Met17_pdr3_int_F and Met17_pdr3_int_R, this fragment serves both as a recombination marker and ligation fragment due to the presence of two *Bam* HI sites in the tails of the forward and reverse primer. The *MET17* fragment was transformed into the *PDR3* locus of strain 1352-Y13363 as described previously (33), hence replacing the *URA3* marker and allowing the use of methionine-controlled promoters and *URA3* based plasmids.

This strain was given the designation HA_SC_1352control. HA_SC_1352control_RED is a fluorescent derivative of HA_SC_1352control, in which an mCherry tag was amplified from plasmid YEP_CHERRY_HIS3 (23) using primer pair Flurotag_marker_F and Flurotag_marker_R and integrated in the *HIS3* locus.

#### HA_SC_Met17_Mpro

An artificial fragment carrying the SARS-COV2 major protease (Mpro, ACCESSION #NC_045512) with an added start and stop codon driven by the *Pichia GAP* promoter and terminated by the *FUM1* terminator (34) was optimized for expression in *S. cerevisiae* and synthesized (GenScript, Piscataway, NJ) then ligated into the *Bam* HI / *Bgl* II site of pPICZ(alpha) A to create plasmid Mpro_pPICZa. Mpro_pPICZa was digested with *Bam* HI and dephosphorylated. Subsequently the above amplified *MET17* fragment was digested with *Bam* HI and ligated to Mpro_pPICZa. A fragment carrying *MET17* ligated to Mpro was amplified from the ligation reaction using primers Mpro_met17_pdr3_int_F and Met17_pdr3_int_R and integrated into the *PDR3* locus of strain 1352-Y13363 to create strain HA_SC_Met17_Mpro (final sequence can be found in Supplementary sequences S1). HA_SC_Met17_Mpro_RED is a fluorescent derivative of HA_SC_Met17_Mpro and was constructed same as HA_SC_1352control_RED. All primers used for strain constructions are listed in Supplementary Table S1.

### Plasmid construction

#### pCM188-GAL1

Empty backbone for galactose driven toxin expression experiments was constructed by ligating the *GAL1* promoter flanked by *Eco* R1 and *Bgl* II into pCM188 (14) digested by *Eco* R1 and *Bam* HI.

#### pCM188-MET3

Empty backbone for methionine-controlled toxin expression experiments was constructed by ligating the *MET3* promoter flanked by *Eco* R1 and *Bam* HI into pCM188 (14) digested by *Eco* R1 and *Bam* HI.

#### PSM (positive selection module)

A chimeric synthetic construct expressing the 41 C-terminal amino acid residues (aa) Leu42 to Trp81 of MazE with an added ATG codon preceding the Leu42 codon, followed by a GGVKLQSGS aa linker sequence containing the Mpro cleavage site VKLQS, joint N-terminally to the full aa sequence of MazF, codon optimized for *S. cerevisiae*, and synthesized (GenScript, Piscataway, NJ). This was then ligated into the *Bam* HI / *Pst* I site of pCM188 to create PSMv3 (full sequence of the MazEF chimera can be found in Supplementary File S2 sequences). PSMv3-GAL was created by ligating the *GAL1* promoter fragment used above to the *Eco* R1 / *Bam* HI sites of PSMv3. PSMv4 was created by ligating the *MET3* promoter fragment used above to *Eco* R1 / *Bam* HI sites of PSMv3.

## Notes

### Competing Interest Statement

The authors have declared no competing interest.

### Summary of Updates

Supplemental files uploaded

