## Supplementary figures and images for "A genetic trap in yeast for inhibitors of SARS-CoV-2 main protease"

### Supplementary Figures

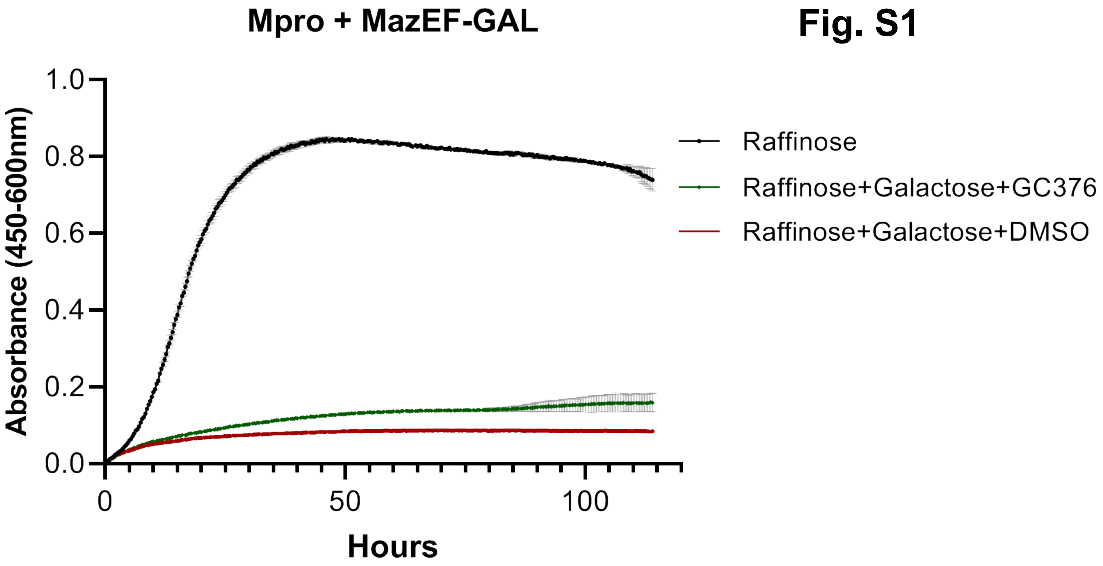

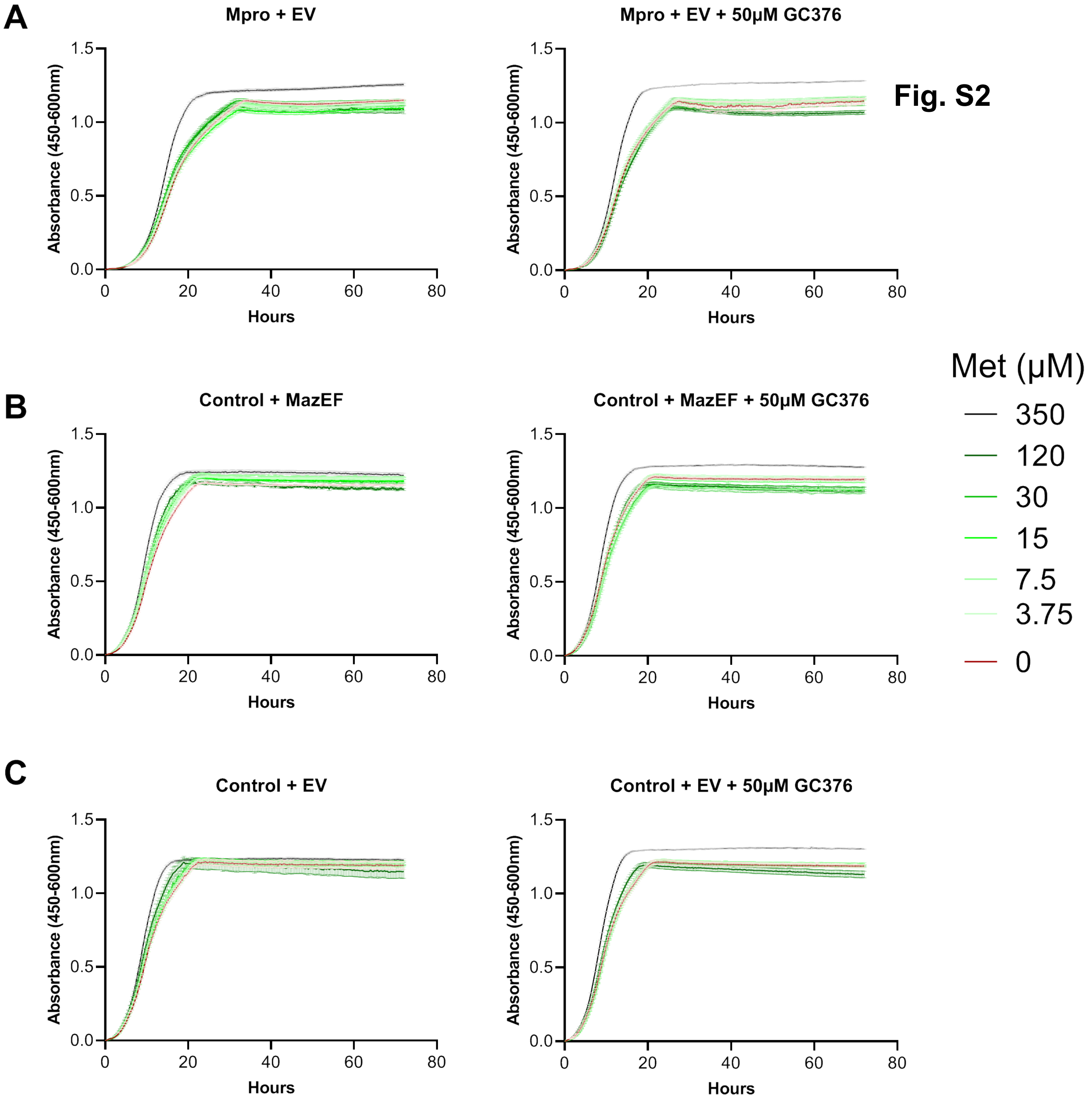

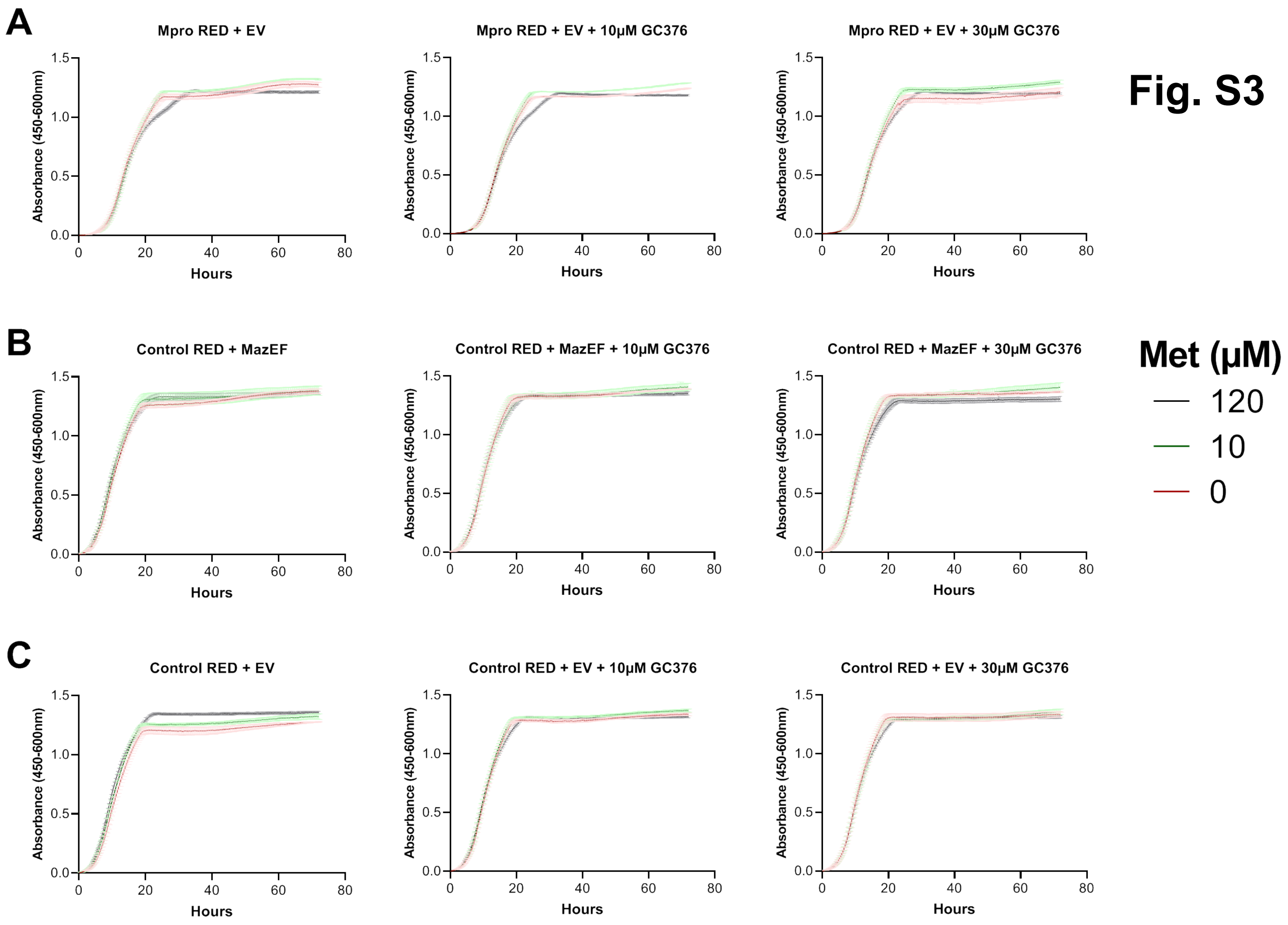

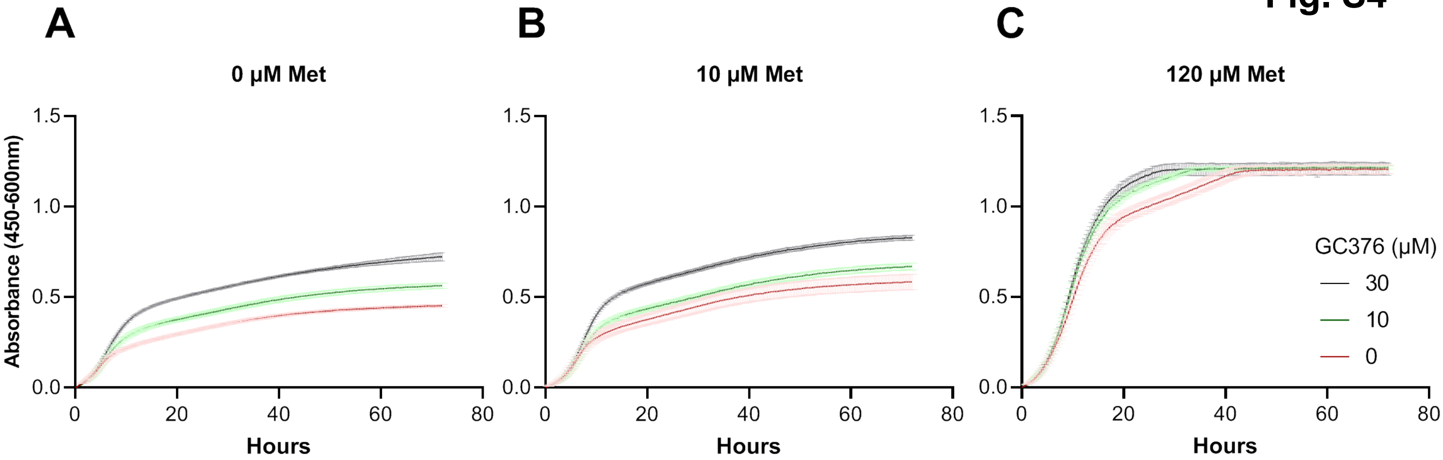

**Fig. S5**

**Mpro vs Control**

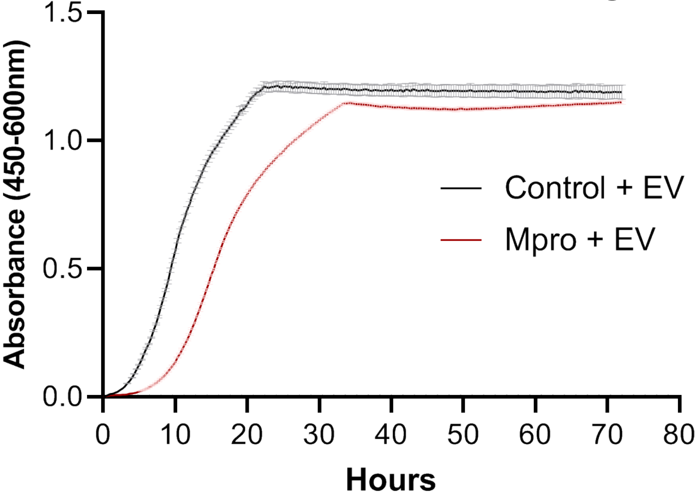
