## Supplementary Files for "A genetic trap in yeast for inhibitors of SARS-CoV-2 main protease"

### Supplementary Table S3. Possible targets for Mpro cleavage in *Saccharomyces cerevisiae*

| Feature Name | Gene Name | HitNu | MatchPattern | MatchSt | MatchS | LocusInfo |
|---|---|---|---|---|---|---|
| YAL053W | FLC2 | 1 | AVLQN | 511 | 515 | Putative calcium channel involved in calcium release under hypotonic stress |
| YBL023C | MCM2 | 1 | TTLQA | 638 | 642 | Protein involved in DNA replication |
| YBL069W | AST1 | 1 | PKLQA | 11 | 15 | Lipid raft associated protein |
| YBR044C | TCM62 | 1 | TRLQS | 97 | 101 | Protein involved in assembly of the succinate dehydrogenase complex |
| YBR047W | FMP23 | 1 | VMLQA | 161 | 165 | Putative protein of unknown function |
| YBR061C | TRM7 | 1 | TTLQA | 94 | 98 | 2'-O-ribose methyltransferase |
| YBR107C | IML3 | 1 | VVLQS | 235 | 239 | Outer kinetochore protein and component of the Ctf19 complex |
| YBR192W | RIM2 | 1 | TRLQS | 78 | 82 | Mitochondrial pyrimidine nucleotide transporter |
| YBR212W | NGR1 | 1 | ATLQS | 180 | 184 | RNA binding protein that negatively regulates growth rate |
| YBR222C | PCS60 | 1 | TKLQS | 120 | 124 | Oxalyl-CoA synthetase |
| YBR236C | ABD1 | 1 | PVLQS | 51 | 55 | Methyltransferase |
| YBR274W | CHK1 | 1 | VVLQS | 66 | 70 | Serine/threonine kinase and DNA damage checkpoint effector |
| YDL024C | DIA3 | 1 | TTLQN | 128 | 132 | Protein of unknown function |
| YDL031W | DBP10 | 1 | AKLQN | 677 | 681 | Putative ATP-dependent RNA helicase of the DEAD-box protein family |
| YDL040C | NAT1 | 1 | ATLQS | 132 | 136 | Subunit of protein N-terminal acetyltransferase NatA |
| YDL159W-A | | 1 | PMLQS | 19 | 23 | Putative protein of unknown function |
| YDR038C | ENA5 | 1 | TTLQS | 1079 | 1083 | Protein with similarity to P-type ATPase sodium pumps |
| YDR158W | HOM2 | 1 | TTLQA | 181 | 185 | Aspartic beta semi-aldehyde dehydrogenase |
| YDR162C | NBP2 | 1 | AVLQN | 29 | 33 | Protein involved in the HOG (high osmolarity glycerol) pathway |
| YDR213W | UPC2 | 1 | ATLQA | 388 | 392 | Sterol regulatory element binding protein |
| YDR252W | BTT1 | 1 | TKLQA | 42 | 46 | Heterotrimeric nascent polypeptide-associated complex beta3 subunit |
| YDR256C | CTA1 | 1 | PVLQA | 346 | 350 | Catalase A |
| YDR268W | MSW1 | 1 | PVLQA | 178 | 182 | Mitochondrial tryptophanyl-tRNA synthetase |
| YDR285W | ZIP1 | 1 | VRLQS | 210 | 214 | Transverse filament protein of the synaptonemal complex |
| YDR295C | HDA2 | 1 | TKLQN | 493 | 497 | Subunit of HDA1 histone deacetylase complex |
| YDR310C | SUM1 | 1 | TMLQN | 172 | 176 | Transcriptional repressor that regulates middle-sporulation genes |
| YDR333C | RQC1 | 1 | AVLQA | 687 | 691 | Component of the ribosome quality control complex (RQC) |
| YDR398W | UTP5 | 1 | PVLQS | 4 | 8 | Subunit of U3-containing Small Subunit (SSU) processome complex |
| YDR440W | DOT1 | 1 | AKLQS | 237 | 241 | Nucleosomal histone H3-Lys79 methylase |
| YDR464W | SPP41 | 1 | AMLQA | 361 | 365 | Protein of unknown function |
| YDR497C | ITR1 | 1 | TMLQN | 502 | 506 | Myo-inositol transporter |
| YEL006W | YEA6 | 1 | TRLQA | 64 | 68 | Mitochondrial NAD+ transporter |
| YEL050C | RML2 | 1 | VRLQS | 287 | 291 | Mitochondrial ribosomal protein of the large subunit (L2) |
| YER006W | NUG1 | 1 | AVLQS | 199 | 203 | GTPase that associates with nuclear 60S pre-ribosomes |
| YER037W | PHM8 | 1 | AKLQS | 213 | 217 | Lysophosphatidic acid (LPA) phosphatase, nucleotidase |
| YER107C | GLE2 | 1 | PTLQA | 308 | 312 | RNA export factor associated with the nuclear pore complex (NPC) |
| YER110C | KAP123 | 1 | AMLQS | 363 | 367 | Karyopherin beta |
| YER141W | COX15 | 1 | VTLQA | 408 | 412 | Heme a synthase |
| YER148W | SPT15 | 1 | PTLQN | 65 | 69 | TATA-binding protein (TBP) |
| YFL022C | FRS2 | 1 | AKLQN | 124 | 128 | Alpha subunit of cytoplasmic phenylalanyl-tRNA synthetase |
| YFR010W | UBP6 | 1 | ATLQA | 122 | 126 | Ubiquitin-specific protease |
| YFR030W | MET10 | 1 | TVLQA | 731 | 735 | Subunit alpha of assimilatory sulfite reductase |
| YFR032C | RRT5 | 1 | VTLQS | 187 | 191 | Putative protein of unknown function |
| YFR041C | ERJ5 | 1 | PKLQN | 52 | 56 | Type I membrane protein with a J domain |
| YGL045W | RIM8 | 1 | VVLQS | 211 | 215 | Protein involved in proteolytic activation of Rim101p |
| YGL073W | HSF1 | 1 | ARLQN | 642 | 646 | Trimeric heat shock transcription factor |
| YGL086W | MAD1 | 1 | TTLQN | 147 | 151 | Coiled-coil protein involved in spindle-assembly checkpoint |
| YGL234W | ADE5,7 | 1 | ARLQS | 59 | 63 | Enzyme of the 'de novo' purine nucleotide biosynthetic pathway |
| YGR130C | | 1 | TRLQA | 742 | 746 | Component of the eisosome with unknown function |
| YGR138C | TPO2 | 1 | ATLQS | 92 | 96 | Polyamine transporter of the major facilitator superfamily |
| YGR162W | TIF4631 | 1 | AKLQS | 152 | 156 | Translation initiation factor eIF4G and scaffold protein |
| YGR250C | RIE1 | 1 | TVLQA | 147 | 151 | RNA binding protein and negative regulator of START |
| YGR251W | NOP19 | 1 | AKLQS | 15 | 19 | Ribosome biogenesis factor |
| YGR257C | MTM1 | 1 | TKLQS | 184 | 188 | Mitochondrial protein of the mitochondrial carrier family |
| YHR006W | STP2 | 1 | AKLQN | 134 | 138 | Transcription factor |
| YHR099W | TRA1 | 1 | ATLQS | 19 | 23 | Subunit of SAGA and NuA4 histone acetyltransferase complexes |
| YHR134W | WSS1 | 1 | AVLQS | 33 | 37 | SUMO-ligase and SUMO-targeted metalloprotease |
| YHR158C | KEL1 | 1 | VRLQS | 833 | 837 | Protein required for proper cell fusion and cell morphology |
| YIL006W | YIA6 | 1 | TRLQA | 103 | 107 | Mitochondrial NAD+ transporter |
| YIL095W | PRK1 | 1 | TRLQN | 122 | 126 | Ser/Thr protein kinase |
| YIL114C | POR2 | 1 | ATLQS | 205 | 209 | Putative mitochondrial porin (voltage-dependent anion channel) |
| YIL130W | ASG1 | 1 | TRLQS | 124 | 128 | Zinc cluster protein proposed to be a transcriptional regulator |
| YIL131C | FKH1 | 1 | TVLQS | 151 | 155 | Forkhead family transcription factor |
| YIR010W | DSN1 | 1 | ATLQS | 41 | 45 | Essential component of the outer kinetochore MIND complex |
| YIR039C | YPS6 | 1 | PMLQA | 281 | 285 | Putative GPI-anchored aspartic protease |
| YJL059W | YHC3 | 1 | TVLQS | 318 | 322 | Protein required for the ATP-dependent transport of arginine |
| YJL122W | ALB1 | 1 | VTLQN | 72 | 76 | Shuttling pre-60S factor |
| YJL132W | | 1 | TKLQA | 21 | 25 | Putative protein of unknown function |
| YJL163C | | 1 | PTLQS | 474 | 478 | Putative protein of unknown function |
| YJL165C | HAL5 | 1 | PVLQA | 73 | 77 | Snf1p-related nutrient-responsive protein kinase |
| YJR045C | SSC1 | 1 | TRLQS | 21 | 25 | Hsp70 family ATPase |

| | | | | | | |
|---|---|---|---|---|---|---|
| YKL041W | VPS24 | 1 | TVLQN | 39 | 43 | One of four subunits of the ESCRT-III complex |
| YKL078W | DHR2 | 1 | ATLQA | 237 | 241 | Predominantly nucleolar DEAH-box ATP-dependent RNA helicase |
| YKL082C | RRP14 | 1 | AMLQA | 331 | 335 | Essential protein, constituent of 66S pre-ribosomal particles |
| YKL120W | OAC1 | 1 | TRLQS | 154 | 158 | Mitochondrial inner membrane transporter |
| YKR009C | FOX2 | 1 | ATLQA | 605 | 609 | 3-hydroxyacyl-CoA dehydrogenase and enoyl-CoA hydratase |
| YKR071C | DRE2 | 1 | TKLQS | 141 | 145 | Component of the cytosolic Fe-S protein assembly (CIA) machinery |
| YKR093W | PTR2 | 1 | AVLQS | 439 | 443 | Integral membrane peptide transporter |
| YLL043W | FPS1 | 1 | AMLQA | 435 | 439 | Aquaglyceroporin, plasma membrane channel |
| YLR007W | NSE1 | 1 | AVLQS | 119 | 123 | Component of the SMC5-SMC6 complex |
| YLR018C | POM34 | 1 | TKLQS | 235 | 239 | Subunit of the transmembrane ring of the nuclear pore complex (NPC) |
| YLR022C | SDO1 | 1 | AKLQA | 150 | 154 | Guanine nucleotide exchange factor (GEF) for Ria1p |
| YLR024C | UBR2 | 1 | TVLQS | 314 | 318 | Cytoplasmic ubiquitin-protein ligase (E3) |
| YLR067C | PET309 | 1 | AVLQS | 382 | 386 | Specific translational activator for the COX1 mRNA |
| YLR117C | CLF1 | 2 | AKLQS | 568 | 572 | Member of the NineTeen Complex (NTC) |
| YLR117C | CLF1 | 2 | VRLQN | 645 | 649 | Member of the NineTeen Complex (NTC) |
| YLR139C | SLS1 | 1 | ARLQS | 341 | 345 | Mitochondrial membrane protein |
| YLR238W | FAR10 | 1 | VKLQN | 333 | 337 | Protein involved in recovery from arrest in response to pheromone |
| YLR320W | MMS22 | 1 | TTLQS | 1421 | 1425 | Subunit of E3 ubiquitin ligase complex involved in replication repair |
| YLR348C | DIC1 | 1 | VRLQA | 39 | 43 | Mitochondrial dicarboxylate carrier |
| YLR381W | CTF3 | 1 | PVLQS | 104 | 108 | Outer kinetochore protein that forms a complex with Mcm16p and Mcm22p |
| YLR384C | IKI3 | 1 | VVLQA | 727 | 731 | Subunit of Elongator complex |
| YLR398C | SKI2 | 1 | TRLQS | 798 | 802 | Ski complex component and putative RNA helicase |
| YLR422W | DCK1 | 1 | TRLQS | 1528 | 1532 | Dock family protein (Dedicator Of CytoKinesis), homolog of human DOCK1 |
| YML104C | MDM1 | 1 | AMLQN | 1100 | 1104 | PtdIns-3-P binding protein that tethers the ER to vacuoles at NVJs |
| YMR039C | SUB1 | 1 | PTLQA | 247 | 251 | Transcriptional regulator |
| YMR126C | DLT1 | 1 | AVLQN | 100 | 104 | Protein of unknown function |
| YMR137C | PSO2 | 1 | TVLQN | 475 | 479 | Nuclease required for DNA single- and double-strand break repair |
| YMR154C | RIM13 | 1 | AKLQN | 107 | 111 | Calpain-like cysteine protease |
| YMR203W | TOM40 | 1 | TTLQA | 273 | 277 | Component of the TOM (translocase of outer membrane) complex |
| YMR258C | ROY1 | 1 | TVLQS | 392 | 396 | GTPase inhibitor with similarity to F-box proteins |
| YMR264W | CUE1 | 1 | TKLQS | 190 | 194 | Ubiquitin-binding protein |
| YNL003C | PET8 | 1 | TRLQA | 30 | 34 | S-adenosylmethionine transporter of the mitochondrial inner membrane |
| YNL020C | ARK1 | 1 | TRLQN | 123 | 127 | Ser/Thr protein kinase |
| YNL085W | MKT1 | 1 | TTLQN | 302 | 306 | Protein similar to nucleases that forms a complex with Pbp1p |
| YNL100W | MIC27 | 1 | VVLQN | 22 | 26 | Component of the MICOS complex |
| YNL123W | NMA111 | 1 | AVLQA | 886 | 890 | Serine protease and general molecular chaperone |
| YNL225C | CNM67 | 1 | TTLQN | 484 | 488 | Component of the spindle pole body outer plaque |
| YNR055C | HOL1 | 1 | TTLQN | 265 | 269 | Putative transporter in the major facilitator superfamily |
| YNR063W | PUL4 | 1 | ATLQS | 407 | 411 | Putative zinc-cluster transcription factor |
| YOL021C | DIS3 | 1 | VVLQA | 94 | 98 | Exosome core complex catalytic subunit |
| YOL031C | SIL1 | 1 | VVLQN | 267 | 271 | Nucleotide exchange factor for the ER lumenal Hsp70 chaperone Kar2p |
| YOL078W | AVO1 | 1 | TRLQN | 1162 | 1166 | Component of a membrane-bound complex containing the Tor2p kinase |
| YOL103W | ITR2 | 1 | TMLQN | 525 | 529 | Myo-inositol transporter |
| YOL140W | ARG8 | 1 | TKLQS | 193 | 197 | Acetylornithine aminotransferase |
| YOR083W | WHI5 | 1 | VKLQN | 191 | 195 | Repressor of G1 transcription |
| YOR100C | CRC1 | 1 | TKLQA | 265 | 269 | Mitochondrial inner membrane carnitine transporter |
| YOR109W | INP53 | 1 | TKLQS | 551 | 555 | Polyphosphatidylinositol phosphatase |
| YOR127W | RGA1 | 1 | PVLQN | 751 | 755 | GTPase-activating protein for polarity-regulator Cdc42p (RhoGAP) |
| YOR156C | NFI1 | 1 | AVLQN | 589 | 593 | SUMO E3 ligase |
| YOR180C | DCI1 | 1 | TVLQS | 51 | 55 | Peroxisomal protein |
| YOR194C | TOA1 | 1 | VKLQA | 200 | 204 | TFIIA large subunit |
| YOR270C | VPH1 | 1 | ATLQA | 347 | 351 | Subunit a of the vacuolar-ATPase V0 domain |
| YPL029W | SUV3 | 1 | PKLQA | 127 | 131 | ATP-dependent RNA helicase |
| YPL037C | EGD1 | 1 | TKLQS | 42 | 46 | Subunit beta1 of the nascent polypeptide-associated complex (NAC) |
| YPL051W | ARL3 | 1 | TTLQS | 112 | 116 | ARF-like small GTPase of the RAS superfamily |
| YPL093W | NOG1 | 1 | AKLQA | 451 | 455 | Putative GTPase |
| YPL166W | ATG29 | 1 | PTLQN | 114 | 118 | Autophagy-specific protein |
| YPL228W | CET1 | 1 | ATLQS | 213 | 217 | RNA 5'-triphosphatase involved in mRNA 5' capping |
| YPL242C | IQG1 | 1 | TKLQS | 543 | 547 | Actin filament binding protein that enhances actin ring formation |
| YPL253C | VIK1 | 2 | ATLQA | 161 | 165 | Protein that forms a kinesin-14 heterodimeric motor with Kar3p |
| YPL253C | VIK1 | 2 | ATLQS | 271 | 275 | Protein that forms a kinesin-14 heterodimeric motor with Kar3p |
| YPR010C | RPA135 | 1 | VMLQS | 163 | 167 | RNA polymerase I second largest subunit A135 |
| YPR046W | MCM16 | 1 | TKLQN | 84 | 88 | Component of the Ctf19 complex and the COMA subcomplex |
| YPR128C | ANT1 | 1 | AMLQS | 248 | 252 | Peroxisomal adenine nucleotide transporter |
| YPR169W | JIP5 | 1 | AKLQS | 57 | 61 | Protein required for biogenesis of the large ribosomal subunit |

**Supplementary File S2**

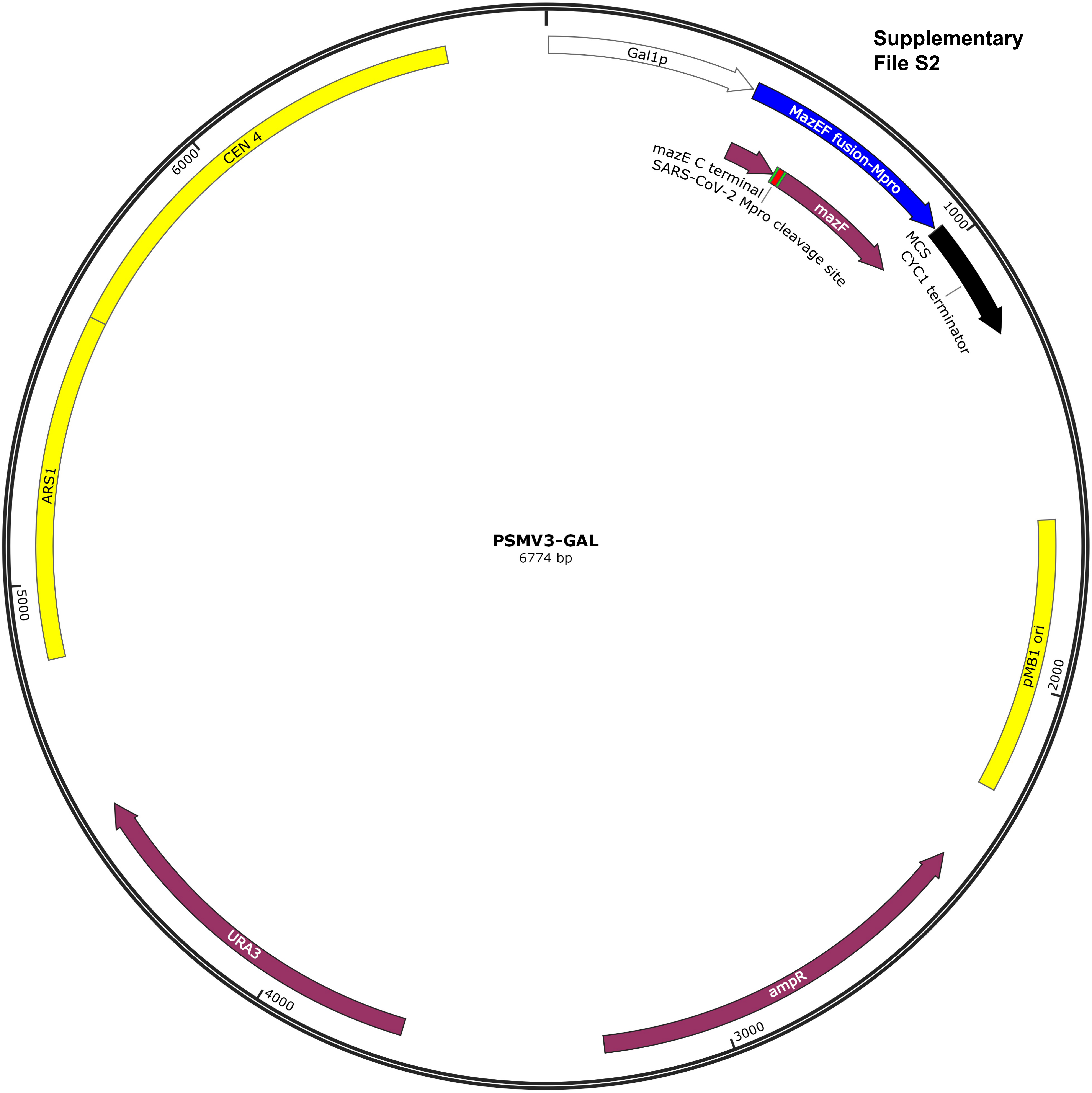

**Supplementary File S3**

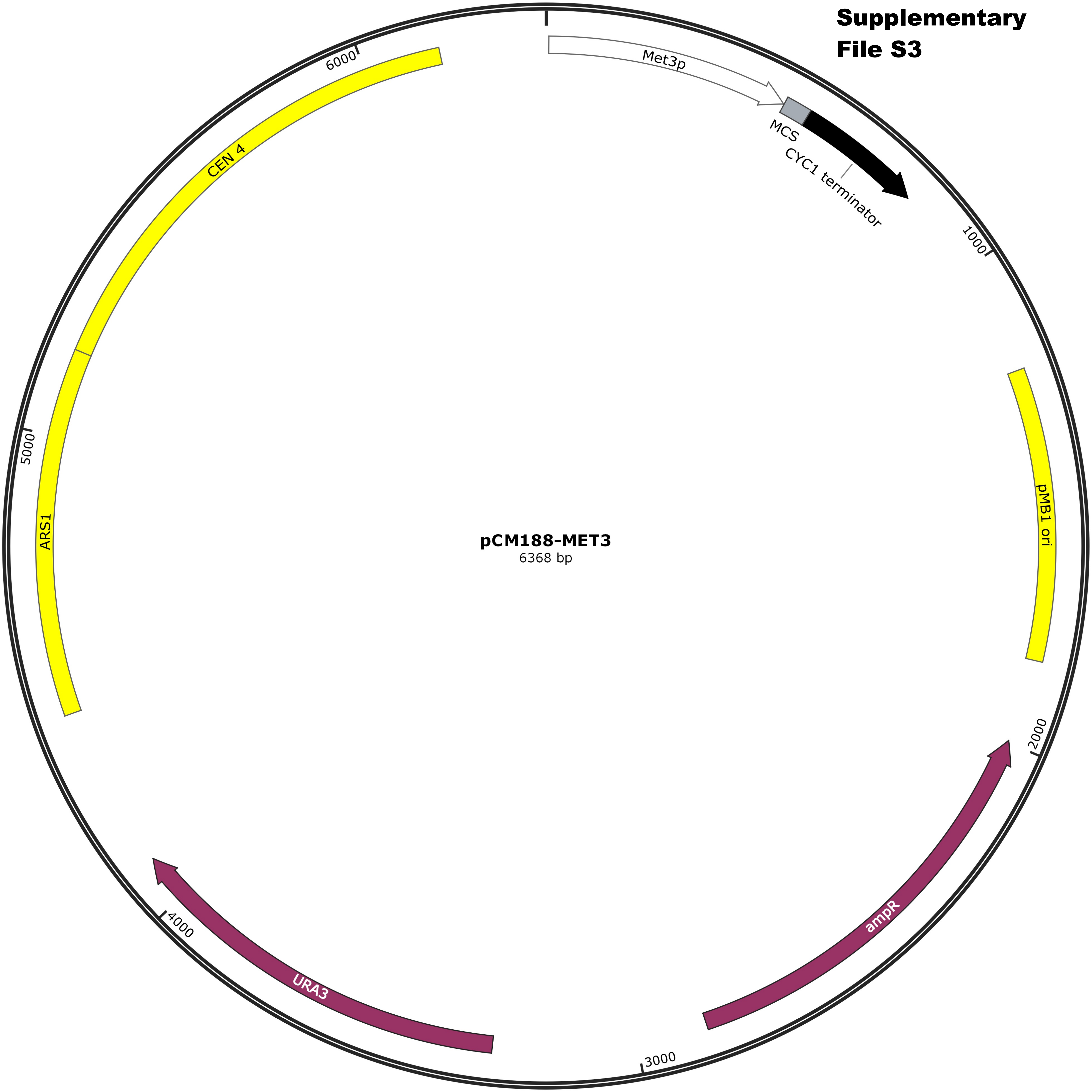

**Supplementary File S4**

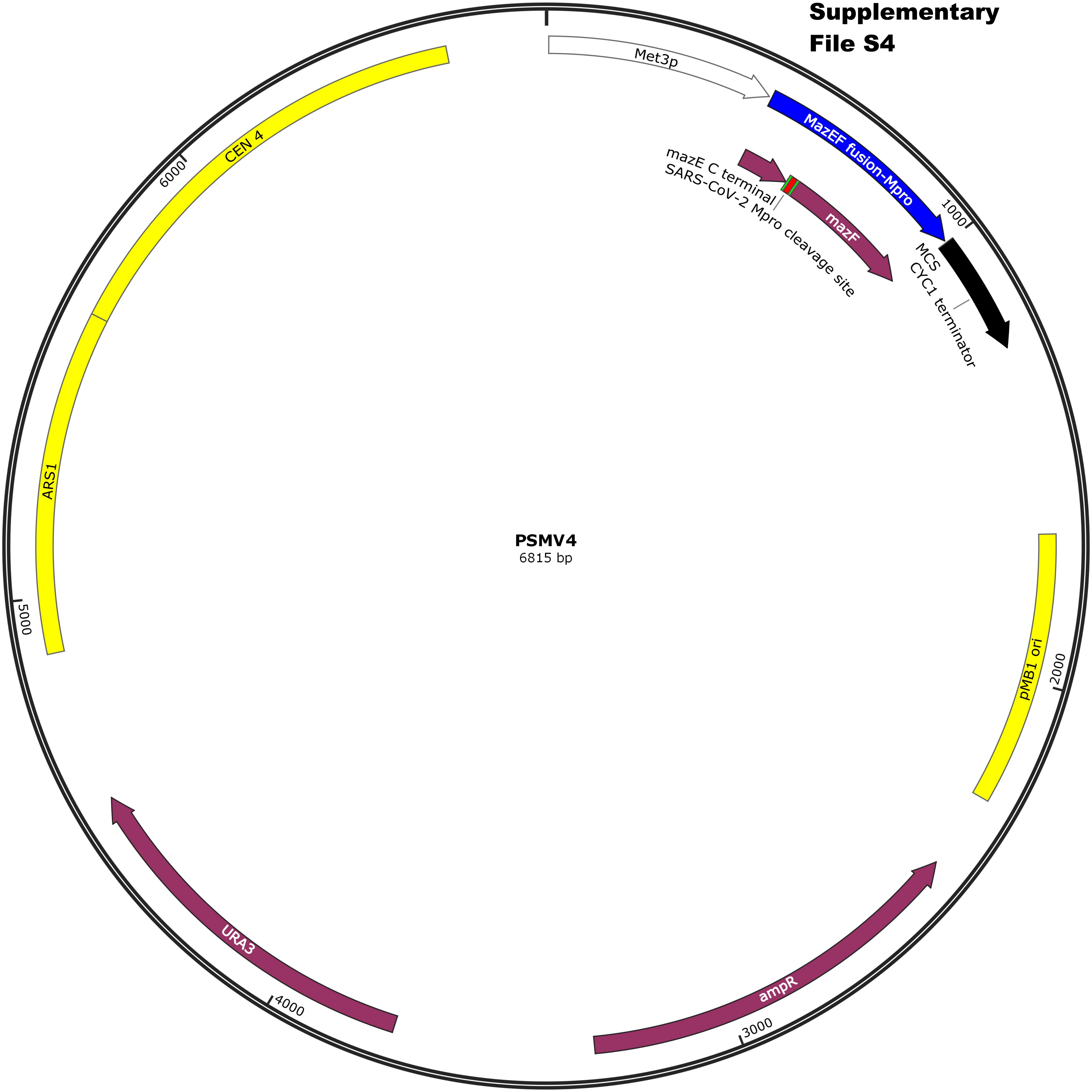
